# Building a Knowledge Graph to Enable Precision Medicine

**DOI:** 10.1101/2022.05.01.489928

**Authors:** Payal Chandak, Kexin Huang, Marinka Zitnik

## Abstract

Developing personalized diagnostic strategies and targeted treatments requires a deep understanding of disease biology and the ability to dissect the relationship between molecular and genetic factors and their phenotypic consequences. However, such knowledge is fragmented across publications, non-standardized research repositories, and evolving ontologies describing various scales of biological organization between genotypes and clinical phenotypes. Here, we present PrimeKG, a precision medicine-oriented knowledge graph that provides a holistic view of diseases. PrimeKG integrates 20 high-quality resources to describe 17,080 diseases with 4,050,249 relationships representing ten major biological scales, including disease-associated protein perturbations, biological processes and pathways, anatomical and phenotypic scales, and the entire range of approved and experimental drugs with their therapeutic action, considerably expanding previous efforts in disease-rooted knowledge graphs. In addition, PrimeKG supports artificial intelligence analyses of how drugs might target disease-associated molecular perturbations by containing an abundance of ‘indications’, ‘contradictions’, and ‘off-label use’ drug-disease edges lacking in other knowledge graphs. We accompany PrimeKG’s graph structure with text descriptions of clinical guidelines to enable multimodal analyses.

## Background & Summary

Precision medicine takes a personalized approach to disease diagnosis and treatment that accounts for the variability in genetics, environment, and lifestyle across individuals^1^. To be precise, medicine must revolve around data and learn from both biomedical knowledge and health records^2^. Nevertheless, many barriers to linking and efficiently exploiting medical information across healthcare organizations and biological scales slow down the research and development of individualized care^2^. While many have acknowledged the difficulties in linking existing biomedical knowledge to patient-level health records^2–5^, few realize that biomedical knowledge is itself fragmented. Biomedical knowledge about complex diseases comes from different organizational scales, including genomics, transcriptomics, proteomics, molecular functions, intra- and inter-cellular communications, phenotypes, therapeutics, and environmental exposures. For any given disease, information from various organizational scales is scattered across individual publications, non-standardized data repositories, evolving ontologies, and clinical guidelines. Developing networked relationships between these sources can support research in disease-rooted precision medicine.

A resource that comprehensively describes the relationships of diseases to biomedical entities would enable the large-scale, data-driven study of human disease. Understanding the connections between diseases, drugs, phenotypes, and other entities could open the doors for many types of research to leverage recent computational advances, including but not limited to the study of disease phenotyping^6–8^, disease etiology^9^, disease similarity^10^, disease diagnosis^11–13^, disease treatments^14^, drug-disease relationships^15–17^, mechanisms of drug action^18^ and resistance^3^, drug repurposing^19–21^, drug discovery^22,23^, adverse drug events^24,25^, combination drug therapies^26^, and so forth. Many researchers have developed knowledge graphs for individual diseases that have helped advance computational precision medicine within their respective disease area^27–42^. Nevertheless, the costs and extended timelines of these individual efforts demonstrate a need for a systematic data resource that could unify existing biomedical knowledge to enable the investigation of diseases at scale.

While many primary data resources contain information about diseases, consolidating them into a comprehensive, disease-rich, and functional knowledge graph presents three challenges. Firstly, existing approaches to network analysis of diseases require expert review and curation of data in the knowledge graph^29,30,43^. While incredibly detailed, such efforts require substantial manual labor and expensive expert input, making them difficult to scale. Secondly, there lacks a con-sistent representation of diseases across biomedical datasets and clinical guidelines. Rather than have a standardized disease ontology, database developers select the ontology that best suits their function from a multitude of biorepositories^44–54^. Because each set of disease vocabulary was tailored for some to serve a unique purpose, their disease encodings overlap unsystematically and are often in conflict. For instance, ICD codes^50^ are optimized for medical billing whereas MedGen^53^, PhenoDB^51^, and Orphanet^48^ focus on rare and genetic diseases. Moreover, expertly curated disease descriptions in medical knowledge repositories do not follow any naming conventions^48,55^. The lack of standardized disease representations and the multimodal nature of the datasets makes it challenging to harmonize biomedical knowledge at scale. Thirdly, the definition of a ‘unique’ disease remains medically and scientifically ambiguous. For instance, while autism spectrum disorder is considered a medical diagnosis, the condition has many subtypes linked to clinically divergent manifestations^56,57^. Clinically studied disease subtypes often do not correlate clearly with those defined in disease ontologies. Although only three subtypes of autism have be clinically identified^57^, the Unified Medical Language System (UMLS)^46^ describes 192, the Monarch Disease Ontology (MONDO)^44^ describes 37, and Orphanet^48^ describes 6. The challenge in reconciling disease entities is only exacerbated by the variety of synonyms and abbreviations available for any particular disease^58^ and the difficulty in linking structured disease entities to unstructured names in text^59^. Meaningful disease entity resolution across multimodal, non-standardized datasets is critical for developing knowledge graphs that will be useful for downstream precision medicine tasks.

While drug repurposing remains the focus of knowledge graph development^33,37,39,42,60–62^, considerable effort has been devoted to building knowledge graphs from biomedical literature^28,31,40^ and clinical records^29,30,34,63^. For example, the SPOKE network is a seminal effort that linked many heterogeneous biomedical databases to build a disease-centric knowledge graph^38^. Although SPOKE is limited to about 200 diseases and lacks multimodal connections between textual clinical guidelines and tabular molecular data, it has enabled many precision medicine efforts, including Nelson *et al.*^35^ who overlaid individual patient-level information onto SPOKE’s biomedical knowledge. Most recently, an initiative from the White House led to the development of The COVID-19 Open Research Dataset (CORD-19)^64^. CORD-19 was able to empower data-driven medicine during the pandemic by facilitating the development of neural search engines for healthcare workers^65,66^ and provided insights into drug repurposing targets^67^. Collectively, these knowledge graphs have lent themselves to a variety of scientific discoveries^68,69^, methodological innovations^70–72^ and biomedical benchmarking^32,36,73^. Large-scale knowledge graphs have facili-tated fruitful research across various problems faced by the biomedical community. Nevertheless, due to the medical heterogeneity of diseases, the multimodal nature of disease information, and the incompatibility of existing disease biorepositories, knowledge graphs focused on diseases have not achieved the scale or impact of many other efforts in this space.

Here, we present the Precision Medicine Knowledge Graph (PrimeKG), a knowledge graph providing a holistic and multimodal view of diseases. We integrate 20 high-quality resources, biorepositories, and ontologies to curate this knowledge graph. PrimeKG systematically captures information about 17,080 diseases with 4,050,249 relationships representing ten major biological scales, including disease-associated perturbations in the proteome, biological processes and pathways, anatomical and phenotypic scales, and the entire range of approved and experimental drugs together with their therapeutic action, considerably expanding previous efforts in disease-rooted knowledge graph creation. We demonstrate that disease nodes in our multi-relational knowledge graph are densely connected to every other node type, including phenotypes, exposures, and seven others. We tune PrimeKG specifically to support artificial intelligence analyses to better understand how drugs might target disease-associated molecular perturbations by including an abundance of ‘indications’, ‘contradictions’, and ‘off-label use’ drug-disease edges, which are usually missing or sparse in other knowledge graphs. We supplement PrimeKG’s rich graph structure with textual descriptions of clinical guidelines for drug and disease nodes to enable multimodal analyses. Finally, we address the disease entity resolution challenge by improving the correspondence between diseases in PrimeKG and disease subtypes found in the clinic, making PrimeKG-based analyses medically meaningful.

## Methods

The Precision Medicine Knowledge Graph (PrimeKG) is heterogeneous, with 10 types of nodes and 30 types of undirected edges. To develop PrimeKG, we retrieved and collated the 20 primary data resources (detailed in Data Records section) as visualized in Figure 2a, identified relations across these resources as shown in Figures 2b and 2c, harmonized them into a rich, heterogeneous network as illustrated in Figure 2c, and augmented the drug and disease nodes in this network with textual descriptions as depicted in Figure 2d.

**Figure 1:**
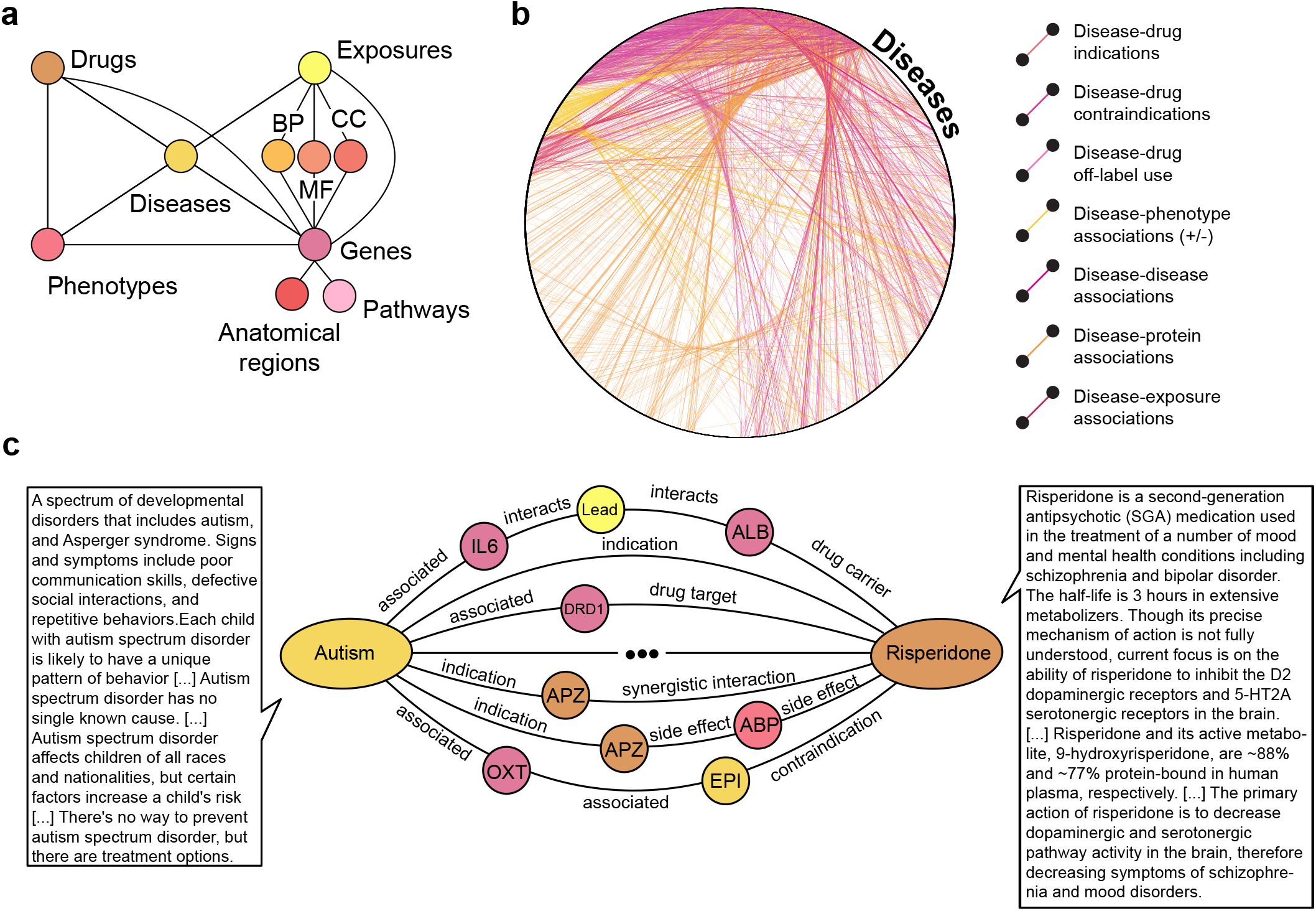
Overview of PrimeKG. **a)** Shown is a schematic overview of the various types of nodes in PrimeKG and the relationships they have with other nodes in the graph. **b)** All disease nodes in PrimeKG shown in a circular layout together with disease-associated information. All relationships between disease nodes and any other node type are depicted here. Disease nodes are densely connected to four other node types in PrimeKG through seven types of relations. **c)** Shown is an example of paths in PrimeKG between the disease node ‘Autism’ and the drug node ‘Risperidone’. Intermediate nodes are colored by their node type from panel a. We also display snippets of text features for both nodes to demonstrate the multimodality of PrimeKG. Abbreviations - MF: molecular function, BP: biological process, CC: cellular component, APZ: Apiprazole, EPI: epilepsy, ABP: abdominal pain, + / - associations: positive and negative associations.

**Figure 2:**
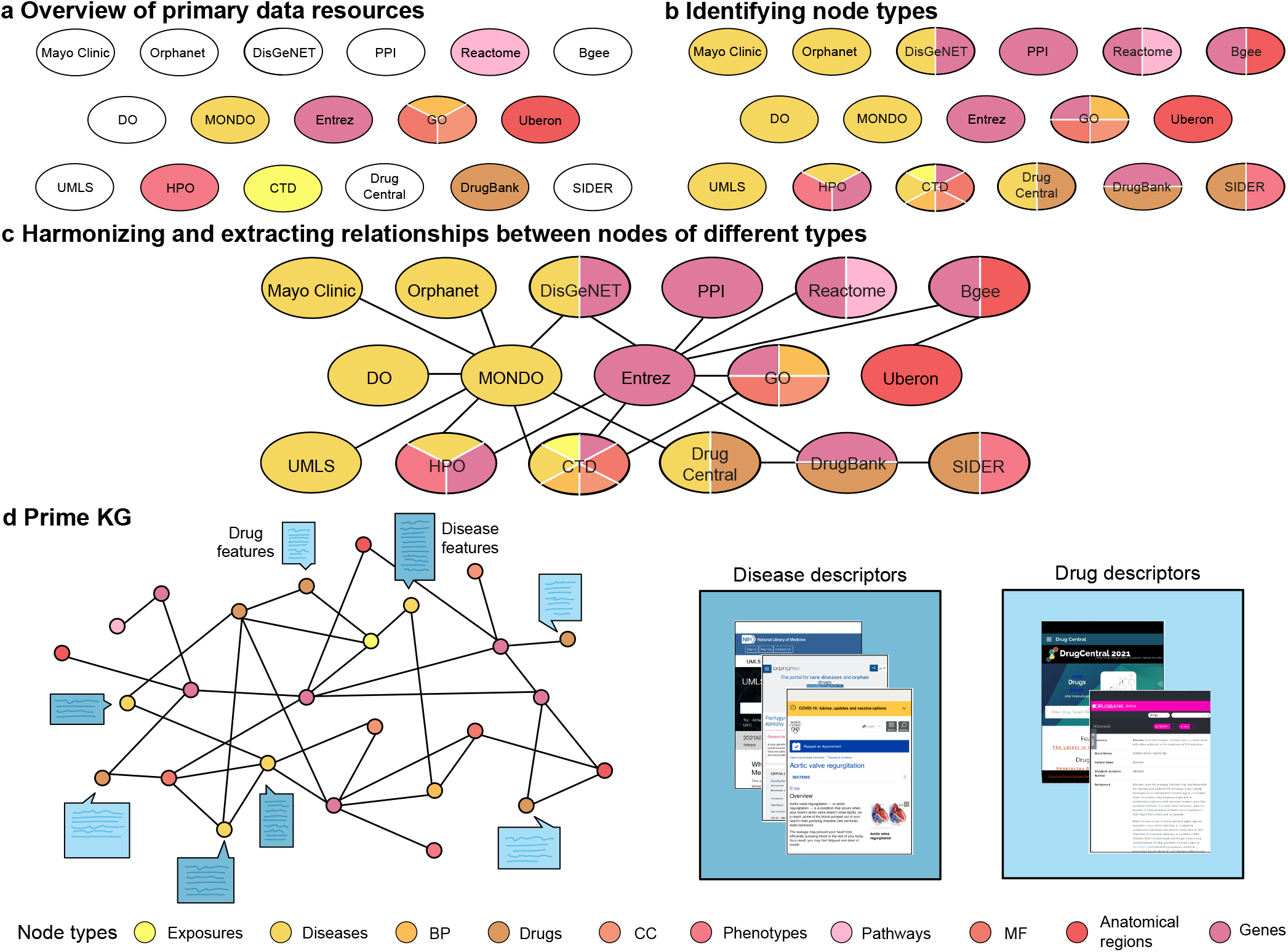
Building PrimeKG. The panels sequentially illustrate the process of developing the Precision Medicine Knowledge Graph. **a)** Shown are 20 primary data resources curated to develop PrimeKG. The colors indicate which database was used to define each node type. **b)** Primary resources are colored by each node type for which they possess information. **c)** Illustrated is the process of harmonizing these primary resources to extract relationships between node types. **d)** The left side illustrates PrimeKG and the right side shows all the textual sources of clinical information on drugs and diseases. The node type legend is consistent across the figure. Abbreviations - MF: molecular function, BP: biological process, CC: cellular component, PPI: protein protein interactions, DO: disease ontology, MONDO: mondo disease ontology, Entrez: entrez gene, GO: gene ontology, UMLS: unified medical language system, HPO: human phenotype ontology, CTD: comparative toxicogenomics database, SIDER: side effect resource.

### A. Curating primary data resources

To develop a comprehensive knowledge graph to study diseases, we considered 20 primary resources and a number of additional repositories of biological and clinical information. Figure 2a provides an overview of all 20 resources. We selected datasets that provided widespread coverage of biomedical entities, including proteins, genes, drugs, diseases, anatomy, biological processes, cellular components, molecular functions, exposures, disease phenotypes and drug side effects. These were high-quality datasets, either expertly curated annotations such as DisGeNet and Mayo Clinic, widely-used standardized ontologies such as Mondo Disease Ontology, or direct readouts of experimental measurements such as Bgee and DrugBank. A complete list of primary resources along with their processing steps is listed in the Data Records section. All our data curation and processing approaches are transparent, fully reproducible, and can be continually adapted as individual data resources evolve and new data become available.

### B. Standardizing and harmonizing data resources

To harmonize these primary data resources into PrimeKG, we selected ontologies for each node type, harmonized datasets into a standardized format, and resolved overlap across ontologies.

#### Defining node types and selecting common ontologies

Our knowledge graph consists of 10 types of nodes. The node types ‘drug’, ‘disease’, ‘anatomy’ and ‘pathway’ are respectively encoded as terms in DrugBank, Mondo, UBERON, and Reactome. Genes and proteins are treated as a single node type, ‘gene/protein’, and identified by Entrez Gene IDs. The node types ‘biological process’, ‘molecular function’, and ‘cellular component’ are defined using Gene Ontology terms. Disease phenotypes extracted from HPO and drug side effects extracted from SIDER are collapsed into a single node type, ‘effect/phenotype’, that is encoded using HPO IDs. Finally, ‘exposure’ nodes are defined using the ExposureStressorID field, which contains MeSH identifiers provided by the Comparative Toxicogenomics Database. This is illustrated in Figure 2a where each database is coloured by the node type it defines. Moving forward, we interchangeably refer to ‘gene/protein’ nodes as proteins and ‘effect/phenotype’ nodes as phenotypes.

#### Harmonizing external data resources

We mapped the aforementioned processed datasets to ensure that all nodes were defined in their respective common ontologies. Next, we identified sources of information across different primary resources for each node type to maximise the number of relationships in PrimeKG (see Figure 2b). We then restructured the datasets to follow the following format. For each node in the knowledge graph, we provide ‘node index’ which is a unique index to identify the node in our KG; ‘node_id’ which indicates the identifier of the node from it’s ontology; ‘node type’ which indicates the node type as defined in our knowledge graph; ‘node name’ which indicates the name of the node as provided by the ontology; and ‘node source’ which indicates the ontology from which ‘node_id’ and ‘node name’ fields were extracted. For each edge in the knowledge graph, we provide ‘relation’ which is the name of the edge type that connects the two nodes; ‘x_index’ which links to the ‘node index’ field; and ‘y_index’ which also links to ‘node_index’. Finally, we renamed columns for consistency, dropped rows with NaN values, dropped duplicated edges and removed self loop edges from each individual dataset.

#### Resolving overlap between phenotype and disease nodes

Since both the Mondo Disease Ontology and Human Phenotype Ontology were developed by the Monarch Initiative, there was considerable overlap between phenotype nodes and disease nodes across the various datasets. Overlapping nodes were defined as effect/phenotype nodes in HPO that (i) had the same ID number as disease nodes in Mondo and (ii) could be mapped from HPO to Mondo using cross-references found in the Mondo ontology. To avoid duplicate nodes, these overlapping phenotype nodes were converted to disease nodes by manipulating edges in various datasets as follows. Let us define the set of overlapping phenotype nodes as *P*. Phenotype-phenotype edges extracted from the HPO ontology were converted to phenotype-disease edges if one phenotype node was in *P* and to disease-disease edges if both phenotype nodes were in *P*. These converted edges were then dropped from the original phenotype-phenotype resource. Protein-phenotype edges extracted from DisGeNet were converted to protein-disease relations if the phenotype node was in *P* and removed from the group of protein-phenotype edges. Finally, for disease-phenotype and drug-phenotype relations, we dropped any edges where the phenotype was in *P*. Adding these edges to drug-disease relations would only introduce unnecessary noise to the indication, contraindication, and off-label use edges.

### C. Building Precision Medicine Knowledge Graph (PrimeKG)

To construct the network structure of PrimeKG, we merged the harmonized primary data resources into a graph and extracted it’s largest connected component as shown in Figure 2c. We integrated the various processed, curated datasets and cleaned the graph by dropping NaN and duplicate edges, adding reverse edges, dropping duplicates again and removing self loops. This version of the knowledge graph is available on our Harvard Dataverse as ‘kg_raw.csv’. To ensure that our knowledge graph was well-connected and did not have any isolated pockets, we extracted its largest connected component. This giant component retained 0.99998% of edges that were present in the original graph. The largest connected component of the knowledge graph is available on our Harvard Dataverse as ‘kg giant.csv’.

### D. Supplementing drug nodes with clinical information

As shown in Figure 2d, we extracted both textual and numerical features for drug nodes in the knowledge graph from DrugBank and Drug Central. Features from DrugBank mapped directly to the knowledge graph since drugs were coded using DrugBank identifiers. Some features had unique attributes for each drug, such as ‘state’, ‘indication’ and ‘mechanism of action’, and others had numerous attributes for each drug, such as ‘group’ and ‘ATC level’. The latter set of features were converted to single text descriptions by joining features using conjunctions such as ‘;’ and ‘and’. Features in Drug Central were mapped to DrugBank IDs using their CAS identifiers, from the vocabulary that was retrieved from DrugBank. Once all features were mapped, text processing involved removing all tokens that are references in DrugBank (for example “[L64839]”) with the help of regular expressions. For the half-life feature, we nullified locations where the text mentioned that no data was available. Finally, we converted numerical features into textual descriptions in order to standardize the feature set.

As an example, let us explore the features available for Prednisolone. Prednisolone is a glucocorticoid similar to cortisol used for its anti-inflammatory, immunosuppressive, anti-neoplastic, and vasoconstrictive effects. Prednisolone has a plasma half life of 2.1-3.5 hours. Prednisolone is indicated to treat endocrine, rheumatic, and hematologic disorders; […] and other conditions like tuberculous meningitis. Corticosteroids binding to the glucocorticoid receptor mediates changes in gene expression that lead to […]. Prednisolone’s protein binding is highly variable, […]. Corticosteroids bind to the glucocorticoid receptor, inhibiting pro-inflammatory signals, and promoting […]. Prednisolone is a solid. Prednisolone is part of Adrenal Cortex Hormones; Adrenals; […] Prednisolone is approved and vet approved. Prednisolone uses Prednisone Action Pathway […] The molecular weight is 360.45. Prednisolone has a topological polar surface area of 94.83. The log p value of is 1.42.

### E. Supplementing disease nodes with clinical information

As shown in Figure 2d, we extracted textual features for diseases nodes in the knowledge graph from the Mondo Disease Ontology, Orphanet, Mayo Clinic, and UMLS. Features from all these sources were mapped to the ‘node id’ field of disease nodes, that was defined using the Mondo Disease Ontology. Since disease nodes were grouped as described in Technical Validation section, many diseases defined in the Mondo Disease Ontology (i.e., many ‘node_id’ values) were collapsed into a single node (i.e., unique ‘node_index’ values). Since disease features are mapped to Mondo identifiers, or the ‘node_id’ field, it is possible for a single disease node in the knowledge graph, defined by a unique ‘node index’, to have multiple feature values for a given feature. We provide the available features in their entirety since we did not have the medical authority to summarize them into single descriptors.

Disease definitions from the Mondo Disease Ontology were directly extracted from the ontology file and unique for each ‘node id’. Disease descriptions extracted from UMLS were mapped from CUI terms to Mondo and as a result, numerous for each ‘node id’. We removed tokens that were references and URLs from UMLS disease descriptions using regular expressions. From Orphanet, we extracted definitions, prevalence, epidemiology, clinical description, and management and treatment. We mapped the features from Orphanet IDs to Mondo, and as a result, there were multiple for each ‘node id’. We used regular expressions to fix formatting errors in the prevalence and epidemiology features.

We extracted the following disease features from Mayo Clinic’s website: symptoms, causes, risk factors, complications, and prevention. Since the Mayo Clinic web-scrapping did not provide a unique identifier in any ontology, we mapped disease names in Mayo Clinic to those in the Mondo Disease Ontology. To develop this mapping, we used a strategy for grouping disease names that is described in detail in the technical validation. Briefly, we conducted automated string matching followed by manual approval of disease name mappings based on their BERT embedding similarity. Automated string matching involved approving exact matches and encapsulated matches, where the name in Mayo was completely present in the name in Mondo. During processing of the symptoms feature, we used regular expressions to extract the end of the text description that explained when to see the doctor as a new and separate feature. Finally, we cleaned the text for formatting errors.

As an example of the depth and breadth of information covered by the disease features, let’s explore Hepatic Porphyria. Per the Mondo Disease Ontology, Hepatic Porphyria is a group of metabolic diseases due to deficiency of one of a number of liver enzymes in the biosynthetic pathway of heme. They are characterized by […]. Clinical features include […]. The UMLS has a very similar disease description. According to Orphanet, it’s a rare sub-group of porphyrias characterized by the occurrence of neuro-visceral attacks with […]. In the majority of European countries, the prevalence of acute hepatic porphyrias is around 1/75000. In 80% of cases the patients are female. All acute hepatic porphyrias can be accompanied by neuro-visceral attacks that appear as […]. The attacks are most commonly triggered by […]. When an acute attack is confirmed, urgent treatment with an injection of […]. According to Mayo Clinic, signs and symptoms of acute porphyria may include: Severe abdominal pain, […], Seizures. All types of porphyria involve a problem in the production of heme […] and a shortage of a specific enzyme determines the type of porphyria. In addition to genetic risks, environmental factors may trigger development of […]. Examples of triggers include: Exposure to sunlight, […]. Possible complications depend on […] During an attack, you may experience […] Although there’s no way to prevent porphyria, if you have the disease, avoid […]. When to see a doctor, […].

## Data Records

We proceed with a detailed description of the 20 primary data resources used to build PrimeKG.

### Bgee gene expression knowledge base in animals

Bgee^74^ contains gene expression patterns across multiple animal species. We retrieved gene expression data for humans from ftp://ftp.bgee.org/current/download/calls/expr_calls/Homo_sapiens_expr_advanced.tsv.gz on 31 May 2021. Processing involved keeping only gold quality calls and ensuring that the anatomical entities were coded using the UBERON ontology. To extract only highly expressed genes in the anatomical entity, we empirically filtered the data to keep data with expression rank less than 25,000. After processing, we had 1,786,311 anatomy-protein associations where gene expression was found to be present or absent.

### Comparative Toxicogenomics Database

The Comparative Toxicogenomics Database (CTD)^75^ is focused on the impact of environmental exposures on human health. We retrieved information about exposures (05/21 version) from http://ctdbase.org/reports/CTD_exposure_events.csv.gz on 9 Jun 2021. Processing involved removing header comments from the csv file. After processing, our data contained 180,976 associations of exposures with proteins, diseases, other exposures, biological processes, molecular functions, and cellular components.

### DisGeNET knowledgebase of gene-disease associations

DisGeNET^76^ is a resource about the relationships between genes and human disease that has been curated by experts. We retrieved curated disease-gene associations (version 7.0) from https://www.disgenet.org/static/disgenet_ap1/files/downloads/curated_gene_disease_associations.tsv.gz on 31 May 2021. The raw data file, ‘curated_gene_disease_associations.tsv’ was not processed further and contains 84,038 associations of genes with diseases, and phenotypes.

### Disease Ontology

Disease Ontology^47^ groups diseases in many meaningful clusters by using clinically relevant characteristics. For instance, diseases are grouped by anatomical entity. We retrieved the ontology from https://raw.githubusercontent.com/DiseaseOntology/HumanDiseaseOntology/main/src/ontology/HumanDO.obo on 29 Jun 2021. The raw data ‘HumanDO.obo’ is mapped to disease nodes in our knowledge graph. Since the Mondo Disease Ontology is not grouped anatomically or by clinical speciality, this will allow users of PrimeKG to explore disease nodes in a medically meaningful format.

### DrugBank

DrugBank^77^ is a resource that contains pharmaceutical knowledge. We retrieved the complete database (version 5.1.8) from https://go.drugbank.com/releases/5-1-8/downloads/all-full-database on 31 May 2021. Processing involved using the beautiful soup package to extract synergistic drug interactions. The processed data contains 2,682,157 associations. We also extracted drug features from the raw data. For over 14,000 drugs, we construct 12 drug features, including group, state, description, mechanism of action, ATC code, pharmacodynamics, half life, protein binding, and pathway.

We also retrieved information about drug targets from https://go.drugbank.com/releases/5-1-8/downloads/target-all-polypeptide-ids, about drug enzymes from https://go.drugbank.com/releases/5-1-8/downloads/enzyme-all-polypeptide-ids, about drug carriers from https://go.drugbank.com/releases/5-1-8/downloads/carrier-all-polypeptide-ids, about drug transporters from https://go.drugbank.com/releases/5-1-8/downloads/transporter-all-polypeptide-ids all on 31 May 2021. Processing involved combining all four resources and mapping gene names from UniProt IDs to NCBI Gene IDs using vocabulary retrieved from HNCG gene names https://www.genenames.org. The processed data contains 26,118 drug-protein interactions.

### Drug Central

Drug Central^78^ is a resource that curates information about drug-disease interactions. We retrieved the Drug Central SQL dump from https://drugcentral.org/ActiveDownload on 1 Jun 2021. The database was loaded into Postgres SQL and drug-disease relationships were extracted. The processed data contains 26,698 indication edges, 8,642 contraindication edges, and 1,917 off-label use edges. We also extracted drug features from the Drug Central SQL dump from the ‘structures’ and ‘structure type’ tables. We extracted features for over 4500 drugs, representing each drug with features including topological polar surface area (TPSA), molecular weight and cLogP. As an example, the features for *Atorvastatin* are: organic structure, molecular weight of 558.65, TPSA of 111.79 and a ClogP value of 4.46.

### Entrez Gene

Entrez Gene^79^ is a resource maintained by the NCBI that contains vast amounts of gene-specific information. We retrieved data about relations between genes and Gene Ontology terms from https://ftp.ncbi.nlm.nih.gov/gene/DATA/gene2go.gz on 31 May 2021. Processing involved using the goatools package^80^ to extract relations between genes and Gene Ontology terms. The processed data contains 297,917 associations of genes with biological processes, molecular functions, and cellular components.

### Gene Ontology

The Gene Ontology^81^ network describes molecular functions, cellular components, and biological processes. We retrieved the ontology from http://purl.obolibrary.org/obo/go/go-basic.obo on 31 May 2021. Processing involved using the goatools package^80^ to extract information for gene ontology terms and relations between go terms. The processed data contains 71,305 hierarchical associations between biological processes, molecular functions, and cellular components.

### Human Phenotype Ontology

The Human Phenotype Ontology^45^ (version hpo-obo@2021-04-13) provides information on phenotypic abnormalities found in diseases. We retrieved the ontology from http://purl.obolibrary.org/obo/hp.obo on 31 May 2021. Processing involved parsing the ontology file to extract phenotype terms in the ontology, parent-child relationships and cross references to other ontologies. The processed data contains disease-phenotype, protein-phenotype, and phenotype-phenotype edges. We also retrieved expertly curated annotations from http://purl.obolibrary.org/obo/hp/hpoa/phenotype.hpoa on 31 May 2021. Additionally, we extracted 218,128 curated positive and negative associations between diseases and phenotypes.

### Mayo Clinic

Mayo Clinic is a nonprofit academic medical center and biomedical research institution focused on integrated health care^55^. On it’s website https://www.mayoclinic.org/diseases-conditions, Mayo Clinic has curated information about symptoms, causes, risk factors, complications and prevention of 2,227 diseases and conditions. We web-scraped this data and extracted descriptions for these diseases and conditions using the *mayo.py* and *diseases.py* scripts on 28 March 2021. The raw data is available at ‘mayo.csv’.

For example, we extracted features of ‘Atrial fibrillation’ from Mayo Clinic. ‘Some people with atrial fibrillation have no symptoms […] others may experience signs and symptoms such as: Palpitations, Weakness, […] and Chest Pain. The disease occurs when ‘the two upper chambers of your heart experience chaotic electrical signals […] As a result, they quiver. The AV node is bombarded with impulses trying to get through to the ventricles’. Certain factors may increase your risk of developing atrial fibrillation including age, heart disease, […] and obesity. Complications include: ‘the chaotic rhythm causing blood to pool in your atria and form clots […] leading to a stroke. […] Atrial fibrillation, especially if not controlled, may weaken the heart and lead to heart failure’. To prevent atrial fibrillation, it’s important to live a heart-healthy lifestyle […] which may include increasing your physical activity, […]. These snippets represent only an overview of over three pages of descriptive features available on Atrial Fibrillation.

### Mondo Disease Ontology

Since the Mondo Disease Ontology^44^ harmonizes diseases from a wide range of ontologies, including OMIM, SNOMED CT, ICD, and MedDRA, it was our preferred ontology for defining diseases. We retrieved the ontology from http://purl.obolibrary.org/obo/mondo.obo on 31 May 2021. Processing involved parsing the ontology file to extract disease terms in the ontology, parent-child relationships, subsets of diseases, cross references to other ontologies, and definitions of disease terms. The processed data contains 64,388 disease-disease edges.

### Orphanet

Orphanet^48^ is a database that focuses on gathering knowledge about rare diseases. The Orphanet website https://www.orpha.net/consor/cgi-bin/Disease_Search_List.php?lng=EN has curated information about definitions, prevalence, management and treatment, epidemiology, and clinical description for 9348 rare diseases. We web-scraped this data and extracted disease features using code available at *orpha.py* on 10 May 2021.

For instance, the rare disease *Hurler syndrome* with Orphanet ID 93473 has the following features. Hurler syndrome is the most severe form of mucopolysaccharidosis type 1, a rare lysosomal storage disease, characterized by skeletal abnormalities, cognitive impairment, heart disease, […] and reduced life expectancy. The prevalence of the Hurler subtype of MPS1 is estimated at 1/200,000 in Europe and one in a million in general. The clinical manifestation of the disease includes ‘ musculoskeletal alterations, cardiomyopathy, […] and neurosensorial hearing loss within the first year of life’. Management of the disease is multidisciplinary: ‘Hematopoietic stem cell transplantation is the treatment of choice as it can prolong survival. […] Enzyme replacement therapy (ERT) with laronidase […] is a lifelong therapy which alleviates non neurological symptoms.’. These descriptions only represent a brief snapshot of the expertly curated knowledge available in Orphanet.

### Four integrated resources of physical protein-protein interactions

Protein-protein interactions are composed of experimentally-verified interactions between proteins. The interactions we consider are diverse in nature and include signalling, regulatory, metabolic-pathway, kinase-substrate and protein complex interactions, which are considered as unweighted and undirected. We use the human PPI network compiled by Menche *et al.*^82^ as the starting resource. This resource integrates several protein-protein interaction databases, including TRANSFAC for regulatory interactions^83^, MINT and IntAct for yeast to hybrid binary interactions^84,85^, and CORUM for protein complex interactions^86^. Additionally, we retrieve protein-protein interaction information from from BioGRID^87^ and STRING^88^ databases. We also consider the human reference interactome (HuRI) generated by Luck *et al*. ^89^, specifically, we use the HI-union, a combination of HuRI and several related efforts to systematically screen for protein-protein interactions. The processed data contains 642,150 edges.

### Reactome pathway database

Reactome^90^ is an open-source, curated database for pathways. We retrieved information about pathways from https://reactome.org/download/current/ReactomePathways.txt, relationships between pathways from https://reactome.org/download/current/ReactomePathwaysRelation.txt and pathway-protein relations from https://reactome.org/download/current/NCBI2Reactome.txt on 31 May 2021. Processing involved extracting ontology information such as hierarchical relationships and extracting pathway-protein interactions. The processed data contains 5,070 pathway-pathway and 85,292 protein-pathway edges.

### Side effect knowledgebases

The Side Effect Resource (SIDER)^91^ contains data about adverse drug reactions. We retrieved side-effect data (SIDER 4.1 version) from http://sideeffects.embl.de/media/download/meddra_all_se.tsv.gz and SIDER’s drug to Anatomical Therapeutic Chemical (ATC) classification mapping from http://sideeffects.embl.de/media/download/drug_atc.tsv on 31 May 2021. Processing involved extracting all side effects where the MedDRA term was coded at the “PT” or preferred term level, and then mapping drugs from STITCH ID to ATC ID. The processed data 202,736 contains drug-phenotype associations.

### Uberon multi-species anatomy ontology

Uberon^92^ is an ontology that contains information about the human anatomy. We retrieved the ontology from http://purl.obolibrary.org/obo/uberon/ext.obo on 31 May 2021. Processing involved extracting information about anatomy nodes and the relationships between them. The processed data 28,064 hierarchical relationships between anatomy nodes.

### UMLS knowledgebase

The Unified Medical Language System (UMLS) Knowledge Source^46^ contains information about biomedical and health related concepts. We retrieved the complete UMLS Metathesauras from https://download.nlm.nih.gov/umls/kss/2021AA/umls-2021AA-metathesaurus.zip on 31 May 2021 in ‘.RRF’ format. To map UMLS CUI terms to the Mondo Disease Ontology, we used the ‘MRCONSO.RRF’ to extract UMLS Concept Unique Identifier (CUI) terms in English. We mapped UMLS CUI terms to Mondo terms in two ways. Firstly, we directly extracted cross references between the two from the Mondo ontology. Secondly, we indirectly mapped UMLS to Mondo using OMIM, NCIT, MESH, MedDRA, ICD 10 and SNOMED CT as intermediate ontologies.

Further, we used ‘MRSTY.RRF’ and ‘MRDEF.RRF’ files to extract definitions for UMLS terms. Of the 127 semantic types present in the ‘MRSTY.RRF’ file, we selected 11 that belonged to the Disorder semantic group in a manner that was consistent with prior work.^93^ These semantic types were Congenital Abnormality, Acquired Abnormality, Injury or Poisoning, Pathologic Function, Disease or Syndrome, Mental or Behavioral Dysfunction, Cell or Molecular Dysfunction, Experimental Model of Disease, Signs and Symptoms, Anatomical Abnormality, and Neoplastic Process. We then used the ‘MRDEF.RRF’ file to extract definitions for CUI terms from sources that were in English.

### Additional vocabularies

We retrieved gene names and mappings between NCBI Entrez IDs and UniProt IDs from https://www.genenames.org/download/custom/ on 31 May 2021. We retrieved the DrugBank drug vocabulary from https://go.drugbank.com/releases/5-1-8/downloads/all-drugbank-vocabulary on 31 May 2021. These were used to map nodes in the knowledge graph to consistent ontologies.

## Technical Validation

As part of the technical validation, we explore the structure and connectivity of PrimeKG.

### Characterizing Precision Medicine Knowledge Graph

PrimeKG contains 129,375 nodes and 8,100,498 edges. Figure 1a shows a schematic overview of the graph structure, containing 10 types of nodes and 30 types of edges. We provide a breakdown of the number of nodes by node type and the number of edges by edge type in Tables 1 and 2, respectively. Figure 1b demonstrates that disease nodes are densely connected to other node types in the knowledge graph. Tables 3 and 4 show statistics on the number of features available for drug and disease nodes. Disease features include information on disease prevalence, symptoms, causes, risk factors, epidemiology, clinical description, management and treatment, complications, prevention, and when to see a doctor. Drug features include molecular weight of chemical compounds, indications, mechanisms of action, pharmacodynamics, protein binding events, and pathway information. This extensive clinical information describing the entire range of drugs and diseases is a unique characteristic of PrimeKG that makes PrimeKG stand out among its peer knowledge graphs. Figure 1c provides an example of the supporting information that is available across these features.

**Table 1:**
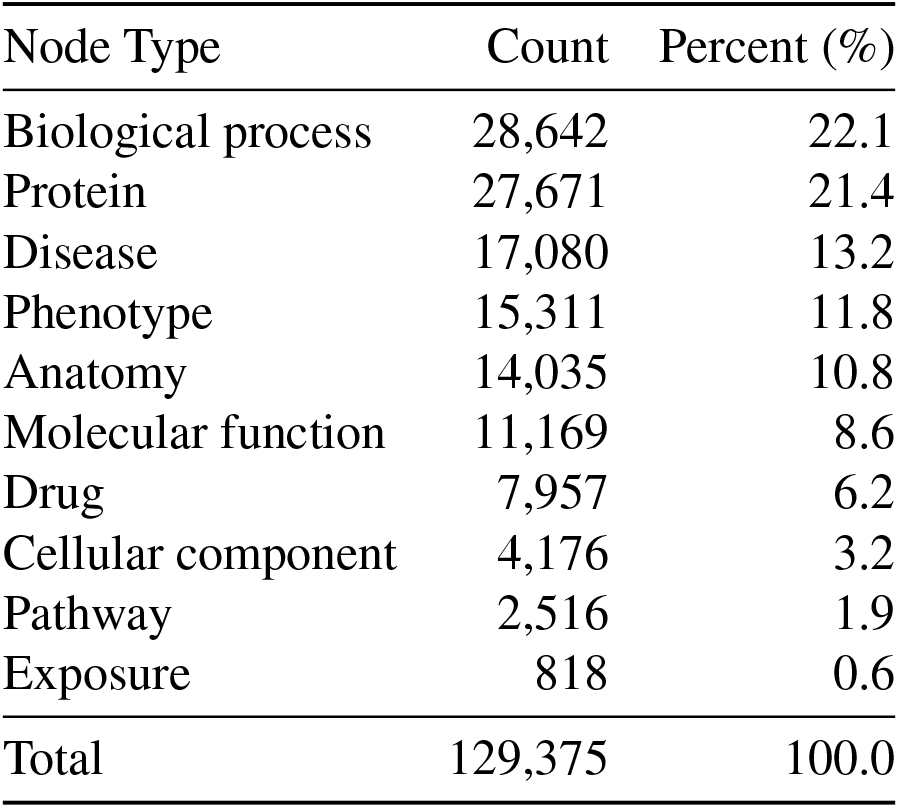
Statistics on nodes in PrimeKG.

**Table 2:** Statistics on edges in PrimeKG.

| Relation type | Count | Percent (%) |
| --- | --- | --- |
| Anatomy - Protein (present) | 3,036,406 | 37.5 |
| Drug - Drug | 2,672,628 | 33.0 |
| Protein - Protein | 642,150 | 7.9 |
| Disease - Phenotype (positive) | 300,634 | 3.7 |
| Biological process - Protein | 289,610 | 3.6 |
| Cellular component - Protein | 166,804 | 2.1 |
| Disease - Protein | 160,822 | 2.0 |
| Molecular function - Protein | 139,060 | 1.7 |
| Drug - Phenotype | 129,568 | 1.6 |
| Biological process - Biological process | 105,772 | 1.3 |
| Pathway - Protein | 85,292 | 1.1 |
| Disease - Disease | 64,388 | 0.8 |
| Drug - Disease (contraindication) | 61,350 | 0.8 |
| Drug - Protein | 51,306 | 0.6 |
| Anatomy - Protein (absent) | 39,774 | 0.5 |
| Phenotype - Phenotype | 37,472 | 0.5 |
| Anatomy - Anatomy | 28,064 | 0.3 |
| Molecular function - Molecular function | 27,148 | 0.3 |
| Drug - Disease (indication) | 18,776 | 0.2 |
| Cellular component - Cellular component | 9,690 | 0.1 |
| Phenotype - Protein | 6,660 | 0.1 |
| Drug - Disease (off-label use) | 5,136 | 0.1 |
| Pathway - Pathway | 5,070 | 0.1 |
| Exposure - Disease | 4,608 | 0.1 |
| Exposure - Exposure | 4,140 | 0.1 |
| Exposure - Biological process | 3,250 | < 0.1 |
| Exposure - Protein | 2,424 | < 0.1 |
| Disease - Phenotype (negative) | 2,386 | < 0.1 |
| Exposure - Molecular function | 90 | < 0.1 |
| Exposure - Cellular component | 20 | < 0.1 |
| Total | 8,100,498 | 100.0 |

**Table 3:** Statistics on drug features in PrimeKG. The count column refers to the number of features including duplicates and the unique column refers to the number of unique features.

| Source | Type of feature | Count | Unique | Percent (%) |
| --- | --- | --- | --- | --- |
| Drug Central<sup>78</sup> | Molecular weight | 2,797 | 2,308 | 35.2 |
|  | TPSA | 2,718 | 2,718 | 34.2 |
|  | cLogP | 2,574 | 980 | 32.3 |
| DrugBank<sup>77</sup> | Group | 7,957 | 7,903 | 100.0 |
|  | State | 6,517 | 6,463 | 81.9 |
|  | Category | 5,431 | 5,431 | 68.3 |
|  | Description | 4,591 | 4,565 | 57.7 |
|  | Indication | 3,393 | 3,076 | 42.6 |
|  | Mechanism of action | 3,242 | 3,161 | 40.7 |
|  | ATC 4 | 2,818 | 1,040 | 35.4 |
|  | ATC 3 | 2,818 | 2,818 | 35.4 |
|  | ATC 2 | 2,818 | 2,818 | 35.4 |
|  | ATC 1 | 2,818 | 2,818 | 35.4 |
|  | Pharmacodynamics | 2,659 | 2,617 | 33.4 |
|  | Half life | 2,063 | 1,893 | 25.9 |
|  | Protein binding | 1,669 | 1,487 | 21.0 |
|  | Pathway | 598 | 598 | 7.5 |

**Table 4:** Statistics on disease features in the knowledge graph. Unprocessed KG refers to the initial knowledge graph assembled from datasets. Processed KG refers to the fully processed PrimeKG, and includes disease groupings. The count column refers to the number of features including duplicates and the unique column refers to the number of unique features.

| Source | Type of feature | Unprocessed KG |  | Processed KG |  |
| --- | --- | --- | --- | --- | --- |
|  |  | Count | Unique | Count | Unique |
| Combined | Combined | 40,068 | 18,152 | 39,800 | 14,252 |
| MONDO Disease Ontology<sup>44</sup> | Definition | 15,238 | 15,238 | 15,238 | 12,001 |
| UMLS<sup>46</sup> | Description | 28,468 | 8,689 | 25,374 | 6,964 |
| Orphanet<sup>48</sup> | Definition | 6,564 | 6,548 | 6,562 | 5,645 |
|  | Prevalence | 3,989 | 3,989 | 3,500 | 3,430 |
|  | Epidemiology | 2,350 | 2,348 | 2,335 | 2,026 |
|  | Clinical description | 2,294 | 2,292 | 2,293 | 1,972 |
|  | Management and treatment | 1,732 | 1,731 | 1,722 | 1,553 |
| Mayo Clinic<sup>55</sup> | Symptoms | 6,642 | 5,789 | 5,140 | 4,470 |
|  | Causes | 6,629 | 5,776 | 5,128 | 4,459 |
|  | Risk factors | 6,284 | 5,501 | 4,898 | 4,299 |
|  | Complications | 5,011 | 4,455 | 3,792 | 3,396 |
|  | Prevention | 2,529 | 2,273 | 1,907 | 1,776 |
|  | When to see a doctor | 5,862 | 5,234 | 4,531 | 4,058 |

### A case study in autism to evaluate the relevance of PrimeKG to clinical presentation of autism

For downstream inferences made using PrimeKG to be conducive to studying human disease, disease nodes in PrimeKG would need to be medically relevant. To this end, we next analyze if PrimeKG’s representation of diseases strongly relates to their clinical presentation by carrying out a case study on autism spectrum disorder. We were motivated to investigate autism because it not only has incredible clinical heterogeneity^94–96^ but this heterogeneity has also been studied to identify clinically meaningfully subtypes^56,57^. We gauged the relevance of disease nodes related to autism in PrimeKG in two steps: first, by performing the entity resolution for autism concepts across all relevant primary data resources (see Methods), and second, by examining the relationship between these autism concepts and clinical subtypes of autism.

We start by exploring whether autism disease nodes in PrimeKG reconciled the variation in autism concepts across databases and ontologies. For example, as demonstrated in Figure 3a, MONDO disease ontology has 37 disease concepts related to autism, whereas the UMLS has 192 autism-associated concepts and Orphanet has 6 autism-associated concepts. Although it is not immediately clear how these concepts relate to each other, we cannot develop a coherent knowledge graph without establishing connections between these concepts. To this end, we overcome this challenge by defining all nodes using the MONDO disease ontology and mapping all other vocabularies to diseases in MONDO as outlined in Figure 3a.

**Figure 3:**
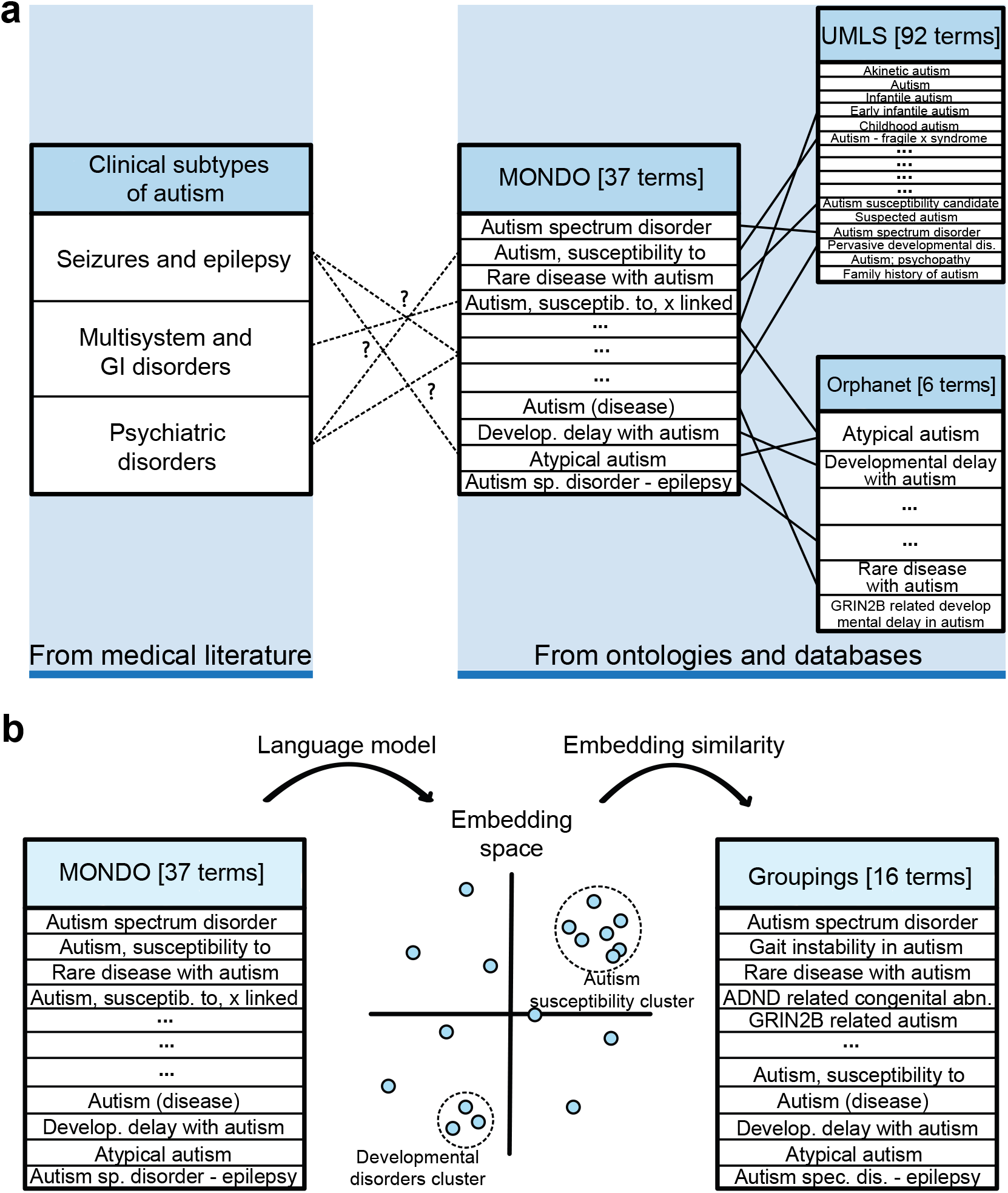
Reconciling autism disease nodes into more medically relevant entities. **a)** The left side shows three clinically determined subtypes of autism. The right side shows autism-related disease terms across three ontologies: MONDO, UMLS, and Orphanet. While we can identify mappings across the ontologies, it is unclear how the terms in any ontology connect to clinical subtypes. **b)** Illustration on how we use a language model, ClinicalBERT, to map terms from MONDO into a latent embedding space. Because the language model can group synonyms in the embedding space, we can cluster MONDO terms with similar semantic and medical meaning by calculating cosine similarity between embeddings of disease concepts. These clusters are created to develop disease groupings as shown on the right in panel b. Abbreviations - MONDO: MONDO disease ontology, UMLS: unified medical language system.

Finally, before using MONDO disease concepts as disease nodes in PrimeKG, we need to assess whether autism disease concepts in MONDO correlate with clinical subtypes of autism. Autism has been shown to manifest as three clinical subgroups characterized primarily by seizures, multisystem and gastrointestinal disorders, and psychiatric disorders^57^. However, it was unclear how the 37 autism disease concepts in MONDO (see Figure 3a) relate to the three clinically defined subtypes. There were many disease concepts in autism, such as ‘Autism, susceptibility to, 1’, ‘Autism, susceptibility to, 2’, ‘Autism, susceptibility to, x-linked’, etc., with no apparent clinical meaning, suggesting that disease nodes in MONDO do not correspond one-to-one to clinical manifestation of autism. For this reason, we developed a strategy to group diseases from MONDO into medically relevant and coherent nodes in PrimeKG. We proceed with describing and evaluating that strategy.

### Computational approaches to grouping disease nodes

As demonstrated in our case study of autism, disease concepts in MONDO may not correlate well with medical subtypes. MONDO contains many repetitive disease entities with no apparent clinical correlation. For this reason, we were motivated to group diseases in MONDO into medically relevant entities. Ideally, we would have preferred to leverage expertise across a wide variety of disease areas when grouping these concepts. However, this approach was time-consuming, expensive, and challenging to execute at scale. Further, disease sub-phenotyping is a relatively new paradigm, and so we anticipated low consensus among medical experts on what constitutes a unique disease.

Since manually grouping diseases with expert supervision was not feasible, we took a semiautomated unsupervised approach to group disease concepts in PrimeKG. Advances in natural language processing, specifically the Bidirectional Encoder Representations from Transformers (BERT) model^97^, allowed us to study similarity between disease concept names. We grouped disease concepts with nearly identical names into a single node with string matching and BERT embedding similarity^97–101^.

We identified disease groups using a string matching strategy across disease names^102^. In this strategy, we selected a disease that ended with a number, or a roman numeral, or any alphanumeric phrase with a length of less than 2, or ‘type’ as the second-last word. Once such a disease was selected, we extracted the primary disease phrase by dropping the ending and used this phrase to find matches. Matches included diseases with the same initial phrase and those containing all phrase words with no other words regardless of word order. For the latter matching criteria, the words ‘type’ and ‘(disease)’ were ignored. In this manner, we grouped disease concepts in MONDO with string matching.

We further tightened groupings identified using string matching by exploring word embedding similarities between disease names, which is visualized in Figure 3b. In natural language processing, word embeddings have been widely and successfully used to resolve conflicting and redundant entities in an unsupervised manner^102–104^, and deep language models such as BERT^97^ can produce semantically meaningful word embeddings. Specifically, ClinicalBERT^105^ is a BERT language model that encodes medical notions of semantics because it has been pre-trained on biomedical knowledge from PubMed^106^ and discharge summaries from MIMIC-III^107^. We used ClinicalBERT to extract word embeddings for disease group names identified during string matching. We also defined similarity between two disease names as the cosine distance between their ClinicalBERT embeddings. Then, after applying an empirically chosen cutoff of similarity ≥ 0.98, we manually approved the suggested disease matches and assigned names to the new groups. Finally, these groupings were applied to the knowledge graph.

After this process, 22,205 disease concepts in MONDO were collapsed into 17,080 grouped diseases, which has resulted in a higher average edge density across diseases and more clinically relevant disease nodes. We anticipate that PrimeKG is a powerful dataset with this grouping because disease representations are concentrated and robust, which, in turn, can enable biological insights gleaned from PrimeKG to be medically relevant.

## Conclusion

The potential uses of PrimeKG are vast. PrimeKG describes drug features on a deeper biological level and disease features on a deeper clinical level, which can be used to explain genotypephenotype associations in terms of genes, pathways, or any other nodes in an extensive knowledge graph, like PrimeKG. Consequently, PrimeKG can be paired with deep graph neural networks^108^ to discover new disease biomarkers, characterize disease processes, hone disease classification, identify phenotypic traits, predict biological mechanisms, and repurpose drugs. With the implementation of machine learning functionality, we anticipate that PrimeKG and similar knowledge graphs will become critical tools in advancing precision medicine.

## Data availability

PrimeKG is hosted on Harvard Dataverse with the following persistent identifier https://doi.org/10.7910/DVN/IXA7BM. We have deposited the knowledge graph along with all relevant intermediate files at this repository.

## Code availability

The PrimeKG’s project website is at https://zitniklab.hms.harvard.edu/projects/PrimeKG. The code to reproduce results, together with documentation and tutorials, is available in PrimeKG’s Github repository at https://github.com/mims-harvard/PrimeKG. In addition, the repository contains information and Python scripts to build new versions of PrimeKG as the underlying primary resources get updated and new data become available.

## Acknowledgements

We would like to thank Bino John, Chris Penland, Nigel Greene, Dominic Williams, and Anna Gogleva for broad discussion on data integration and knowledge graph creation. We also want to thank Jingyi Liu for help with retrieving and processing primary data resources and Michelle M. Li and Emily Alsentzer for helpful discussion on ensuring high-quality of PrimeKG. M.Z. is supported, in part, by NSF under Nos. IIS-2030459 and IIS-2033384, US Air Force Contract No. FA8702-15-D-0001, Harvard Data Science Initiative, Amazon Research Award, Bayer Early Excellence in Science Award, AstraZeneca Research, and Roche Alliance with Distinguished Scientists Award. Any opinions, findings, conclusions or recommendations expressed in this material are those of the authors and do not necessarily reflect the views of the funders.

## Author contributions

P.C., K.H., and M.Z. contributed new analytic tools and wrote the manuscript. P.C. retrieved, processed, harmonized and datasets. P.C. analyzed the resulting knowledge graph. M.Z. designed the study.

## Competing interests

The authors declare no competing interests.

## Notes

https://zitniklab.hms.harvard.edu/projects/PrimeKG/

https://github.com/mims-harvard/PrimeKG

https://doi.org/10.7910/DVN/IXA7BM

### Competing Interest Statement

The authors have declared no competing interest.

### Summary of Updates

Minor edits to finalize the manuscript

